# PIKfyve inhibition reveals a novel role for Inpp4b in the regulation of PtdIns(3)P and lysosome dynamics

**DOI:** 10.1101/2021.12.13.472440

**Authors:** Golam T. Saffi, Emily M. Mangialardi, Jean Vacher, Roberto J. Botelho, Leonardo Salmena

**Affiliations:** Department of Pharmacology & Toxicology, University of Toronto, Toronto, Ontario, Canada; Institut de Recherches Cliniques de Montréal (IRCM), Département de Médecine, Université de Montréal, Montréal, Québec, Canada; Department of Chemistry and Biology, Ryerson University, Toronto, Ontario, Canada; Princess Margaret Cancer Centre, University Health Network, Toronto, Ontario, Canada

**Keywords:** Inpp4b, PIKfyve, Apilimod, Lysosome, PtdIns(3)P

## Abstract

Lysosome membranes contain diverse phosphoinositide (PtdIns) lipids that co-ordinate lysosome function and dynamics. The PtdIns repertoire on lysosomes is tightly regulated by the action of diverse PtdIns kinases and phosphatases. Specific roles for PtdIns in lysosomal function and dynamics are currently unclear and require further investigation. PIKfyve, a lipid kinase which synthesizes PtdIns(3,5)P_2_ from PtdIns(3)P, controls lysosome “fusion-fission” cycles dynamics, autophagosome turnover and endocytic cargo delivery. We have recently characterized a role for INPP4B, a PtdIns phosphatase which hydrolyses PtdIns(3,4)P_2_ to form PtdIns(3)P, in the regulation of lysosomal biogenesis and function. To gain a better understanding of PtdIns homeostasis on lysosomes, we investigated the consequence of disrupting PIKfyve in *Inpp4b*-deficient mouse embryonic fibroblasts. Surprisingly, simultaneous inhibition of Inpp4b and PIKfyve functions impair lysosome “fission” dynamics and thereby exacerbate lysosome enlargement, inhibit autophagic flux. Further examination into the underlying processes that may explain exaggerated lysosome enlargement revealed elevated levels of lysosome associated PtdIns(3)P as contributing factors that control lysosome morphology in cells where Inpp4b and PIKfyve are disrupted. Overall, our study suggests that lysosomal functions are regulated by Inpp4b, through a paradoxical role in suppressing the induction of PtdIns(3)P production.

**SUMMARY STATEMENT:** In this study we identify a novel crosstalk between Inpp4b and PIKfyve. This crosstalk regulates lysosome membrane phosphoinositide composition and lysosomal homeostasis. Our data demonstrate that Inpp4b restricts VPS34-dependent induction of PtdIns(3)P levels, which is activated upon apilimod-mediated PIKfyve inhibition. Through this mechanism, Inpp4b contributes to the regulation of lysosomal membrane dynamics and homeostasis.

## INTRODUCTION

Lysosomes are membrane bound organelles that serve as a cell’s main degradative centre (Luzio et al., 2000; Settembre and Ballabio, 2014). Lysosomes are also key sentinels of cellular nutrient concentration and metabolic activity, and have a critical role in initiating diverse signal transduction pathways (Savini et al., 2019; Lamming and Bar-Peled, 2019; Inpanathan and Botelho, 2019). Lysosomal membranes have been demonstrated to contain diverse phosphoinositides (PtdIns), known to control lysosome function, dynamics, and homeostasis through the coordinated recruitment of critical proteins (Ebner et al., 2019). PtdIns exist in seven different forms which are defined by the phosphorylation status of the 3, 4, and/or 5 positions of their inositol head group, a process controlled by a number of cellular kinases and phosphatases with PtdIns specificity (Sasaki et al., 2009; Dyson et al., 2012).

Of particular importance at the lysosomal membrane is phosphatidylinositol-3,5-bisphosphate [PtdIns(3,5)P_2_], which is generated through conversion of phosphatidylinositol-3-monophosphate [PtdIns(3)P] by the Phosphoinositide Kinase, FYVE-Type Zinc Finger Containing (PIKfyve) (McCartney et al., 2014; Sbrissa et al., 1999). Pharmacological or genetic inhibition of PIKfyve and subsequent depletion of PtdIns(3,5)P_2_ disrupts lysosomal processes including autophagic flux, endocytic and phagocytic cargo delivery to lysosomes, substrate export from lysosome, and calcium release (Bissig et al., 2017; Sharma et al., 2019; Dayam et al., 2015; Mironova et al., 2016). Additionally, inhibition of PIKfyve leads to a dramatic enlargement of lysosomes, a phenomenon explained in part by the observation of defective membrane recycling from endosomes and/or lysosomes (Ho et al., 2012; McCartney et al., 2014). More recently, Choy and colleagues attributed lysosome enlargement to the disruption of lysosome fusion-fission cycling and/or disruption of “kiss-and-run”, a term which describes a transient membrane fusion event followed by rapid fission event that permits content exchange between lysosomes (Choy et al., 2018; Bright et al., 2005; Duclos et al., 2003; Bissig et al., 2017). In their model, PIKfyve inhibition compromises fission and/or the “run” event. In other words, disrupted fission prom**IN**otes lysosome coalescence, increased individual lysosome volume, and thereby reducing total lysosome numbers (Choy et al., 2018; Saffi and Botelho, 2019).

Emerging evidence suggests that other PtdIns including PtdIns(3)P, PtdIns(4)P and PtdIns(3,4)P_2_ may also have important roles in regulating lysosomal homeostasis. For instance, PtdIns(3)P has been implicated in regulating lysosomal positioning and lysosomal mTORC1 activity (Hong et al., 2017). PtdIns(4)P has demonstrated important functions in lysosomal membrane fusion in late endocytic trafficking (Jeschke and Haas, 2018). PtdIns(3,4)P_2_ was also demonstrated to promote repression of mTORC1 activity and cell growth through functions on lysosomal membranes (Marat et al., 2017). Notably, Inositol polyphosphate 4-phosphatase type II (INPP4B), a PtdIns phosphatase that dephosphorylates PtdIns(3,4)P_2_ to form PtdIns(3)P has been recently demonstrated to have roles in late endosome and lysosome function in diverse cancer settings (Rodgers et al., 2021; Woolley et al., 2021). Expression of the lipid phosphatase INPP4B leads to depleted intracellular PtdIns(3,4)P_2_ and promoted endosomal trafficking of cargo towards lysosomes (Rodgers et al., 2021). Nonetheless, roles for PtdIns(3,4)P_2_ and PtdIns(3)P in lysosomal function and dynamics are currently unclear and require further investigation.

In this study, we investigated PIKfyve inhibition in *Inpp4b^+/+^* and *Inpp4b^-/-^* mouse embryonic fibroblasts (MEF) to gain insight into functional interactions between PIKfyve and INPP4B, and consequently the role of related PtdIns in lysosomal dynamics. Surprisingly, PIKfyve-inhibition in *Inpp4b^-/-^* fibroblasts leads to very massively enlarged lysosomes compared to the enlarged lysosomes produced in *Inpp4b^+/+^* fibroblasts. These findings suggest the existence of coordinated functions for Inpp4b and PIKfyve in maintaining normal lysosomal homeostasis, dynamics and function. Paradoxically, our data point to a novel role for Inpp4b in suppressing VPS34-dependent induction of PtdIns(3)P upon apilimod-mediated PIKfyve inhibition.

## RESULTS

### PIKfyve inhibition in Inpp4b-null MEF leads to formation of massively enlarged LAMP1^+^ vacuoles

PIKfyve-inhibition leads to a plethora of intracellular consequences (Bissig et al., 2017; Sharma et al., 2019; Dayam et al., 2015; Mironova et al., 2016), the most striking of which is the formation of enlarged cellular vacuoles (Choy et al., 2018; McCartney et al., 2014; Ikonomov et al., 2001). Emerging roles for INPP4B in regulating lysosomal funtions have also been reported (Rodgers et al., 2021; Woolley et al., 2021). To gain insight into any functional interactions between PIKfyve and INPP4B, we treated *Inpp4b^+/+^* and *Inpp4b^-/-^* MEF with the specific PIKfyve inhibitor apilimod for 48 hours. Light microscopy revealed the generation of numerous enlarged cellular vacuoles in *Inpp4b^+/+^* MEF; strikingly *Inpp4b^-/-^* MEF developed many more massively enlarged vacuoles (**Fig. 1A**). Immunostaining of endogenous LAMP1 confirmed that all enlarged vacuoles were LAMP1^+^, indicating a lysosomal origin (**Fig. 1B,C**). Quantitation of LAMP1 staining and immunoblotting revealed that total lysosomal content was reduced in *Inpp4b^-/-^* compared to *Inpp4b^+/+^* MEF, in the steady state (**Fig. 1D and 1G**. This was also confirmed by lysotracker staining in *Inpp4b^-/-^* MEF (**Fig. S1A-C**), together suggesting that *Inpp4b*-deficiency leads to decreased total lysosome content.

**Figure 1.**
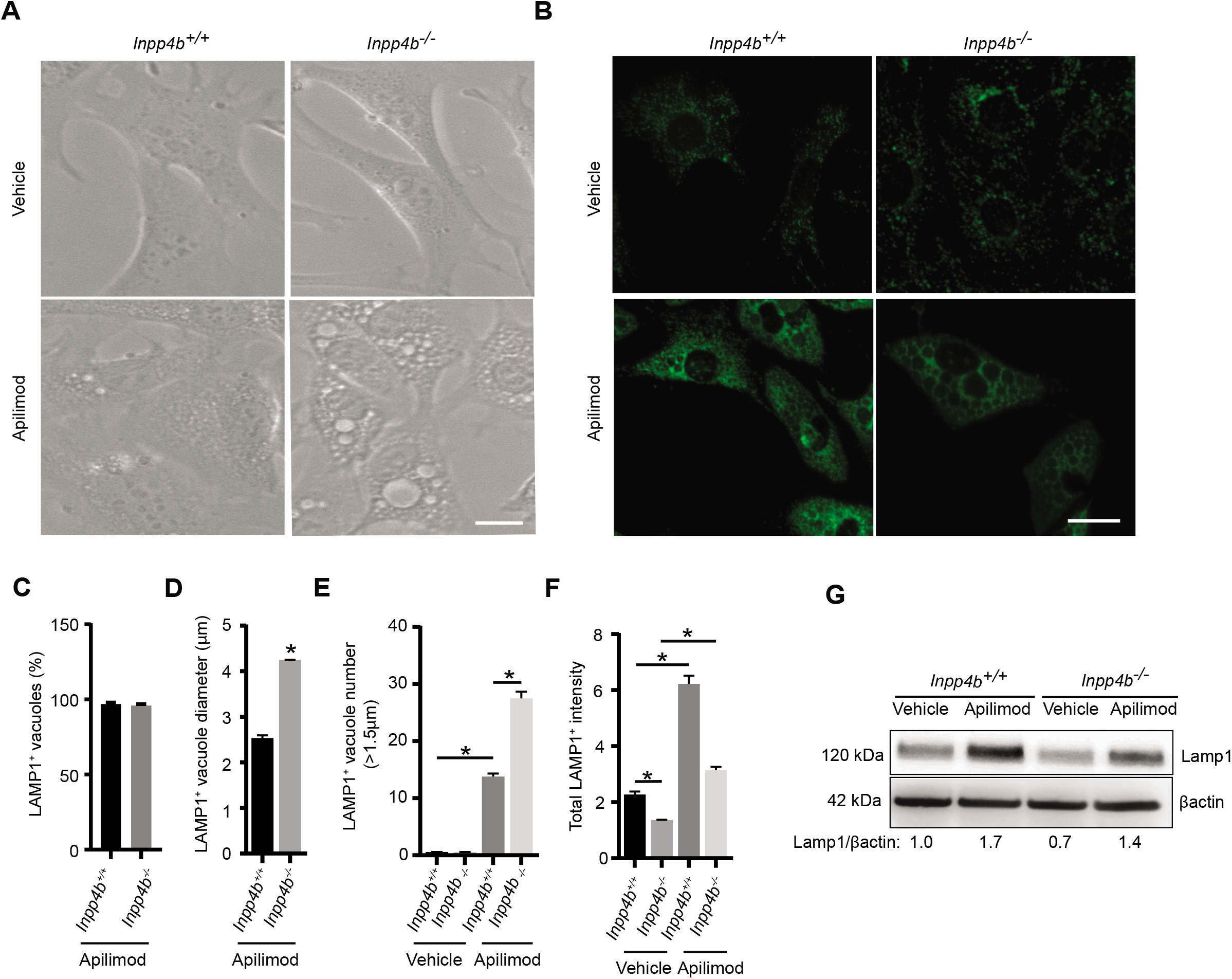
Inpp4b depletion regulates lysosome morphology in response to PIKfyve suppression and reduces lysosome levels. **(A)** *Inpp4b^+/+^* or *Inpp4b^-/-^* MEF cells were incubated with vehicle or 10 nM Apilimod for 48 h followed by imaging with DIC optics. (**B)** Fluorescence micrographs of LAMP1 immunostaining for Inpp4b^+/+^ or Inpp4b^-/-^ MEFs treated with vehicle or 10 nM apilimod for 48 h. (**C**) Percentage of apilimod treated vacuoles positive for LAMP1, (**D**) LAMP1 puncta intensity, (**E**) Mean vacuole diameter (μm) positive for LAMP1 and (**F**) LAMP1 positive vacuole number (>1.5 μm in diameter) per cell across indicated treatments. Scale Bar: 25 μm. (**G**) Representative immunoblot of *Inpp4b^+/+^* or *Inpp4b^-/-^* MEFs treated with vehicle or apilimod 10 nM for 48 h followed by assessment of LAMP1 or □actin as indicated. Data represent ± SEM from three independent experiments with 25-30 cells assessed per treatment condition per experiment for B-F. Statistical significance measured by one way ANOVA and Tukey’s post-hoc represented as * in comparison to indicated conditions (*P* < 0.05).

Apilimod treatment significantly increased total LAMP1 and Lysotracker levels in both *Inpp4b^+/+^* and *Inpp4b^-/-^* MEF in a proportional manner, indicating that apilimod promotes total lysosomal content in MEF (**Fig. 1D, Fig. S1B,C**). Strikingly, *Inpp4b^-/-^* MEF developed significantly more enlarged vacuoles with a significantly larger mean diameter as compared to *Inpp4b^+/+^* MEF (**Fig. 1E,F**). Identical results were observed upon ectopic expression of a mCherry-tagged LAMP1 in *Inpp4b^+/+^* and *Inpp4b^-/-^* MEF (**Fig. S1D-F**).

To further support the lysosomal identity of the enlarged vacuoles formed by apilimod treatment, we stained cells with endosomal markers, EEA1(Wilson et al., 2000) and GFP-2x-FYVE^HRS^ (Chintaluri et al., 2018). EEA1+ vacuoles were spatially separated from LAMP1+ vacuoles, and LAMP1+ vacuoles were generally absent of EEA1 staining, and this segregation was maintained upon apilimod treatment (**Fig. S2A,C**). Likewise, GFP-2x-FYVE^HRS^ structures were absent from vacuoles formed in apilimod-treated *Inpp4b^-/-^* MEF (**Fig. S2B,D**), suggestive of vacuoles with lysosomal identity.

To confirm that these phenotypes in response to apilimod were indeed a result of *Inpp4b*-deficiency, we measured vacuole size and number in apilimod treated MEF that were transiently transfected with constructs that express *EGFP, Inpp4b-EGFP* or catalytically dead *Inpp4b(C845A)-EGFP*. *Inpp4b-EGFP* expression in *Inpp4b^-/-^* MEF was able to significantly reduce the number and size of enlarged vacuoles as compared to untransfected or EGFP control transfected *Inpp4b^-/-^* MEF (**Fig S1G-I**). By contrast, expression of *Inpp4b(C845A)EGFP* in *Inpp4b^-/-^* MEF was unable to alter number and size of enlarged vacuoles (**Fig S1G-I**), suggesting that active Inpp4b phosphatase can limit enlargement upon apilimod treatment. Moreover, exacerbated lysosomal enlargement in INPP4B-deficiency upon was generalizable to human cells, as measured in U2OS cells upon INPP4B knockdown with RNAi (**Fig. S3A-E**). Overall, apilimod treatment revealed that *Inpp4b^-/-^* MEF demonstrate an exacerbated lysosomal vacuolation phenotype compared to *Inpp4b^+/+^* MEF that can be rescued by *wild-type* Inpp4b, but not a catalytically dead-Inpp4b. These data suggest an important role for Inpp4b phosphatase function in maintaining lysosomal dynamics and homeostasis.

### Inpp4b-deficiency exacerbates apilimod-mediated inhibition of endocytic trafficking to lysosomes

We attempted to measure lysosome dynamics using Lucifer yellow (LY), a membrane impermeable fluorescent dye that is internalized by endocytosis and accumulates within lysosomes. LY fluorescence provides an effective tool to specifically delineate lysosomes for accurate measurement of lysosome number and volume (Chaurra et al., 2011; Baluska et al., 2004). In this experiment, MEF were treated with vehicle or apilimod for 48 hours prior to LY pulses of 1, 2 and 4 hours. First, total intracellular LY-fluorescence as measured by flow cytometry demonstrated that apilimod results in the accumulation of LY, however both *Inpp4b^+/+^* and *Inpp4b^-/-^* MEF demonstrate similar levels of LY accumulation for up to 4 hours (**Fig. S4A**). To monitor LY trafficking to lysosomes, ectopic expression of mCherry-LAMP1 was used to label lysosomes (**Fig. S4B**). Surprisingly, apilimod treatment demonstrated a markedly reduced accumulation of LY to lysosomes; comparatively *Inpp4b^-/-^* MEF has significantly less LY accumulation at lysosomes compared to *Inpp4b^+/+^* MEF (**Fig. S4B,C**). To measure if the apilimod-mediated reduced accumulation of cargo in lysosomes also affects degradation, we performed the same assay but replaced LY with DQ-BSA, bovine serum albumin labelled with a self-quenched BODIPY TR-X dye which is cleaved by lysosomal hydrolases, resulting in bright green fluorescent signal (Marwaha and Sharma, 2017; Gray et al., 2016). Fluorescence activation of DQ-BSA was delayed but not inhibited in apilimod treated cells, (**Fig. S4D**), suggesting that degradation of lysosomal cargo was functional. These findings suggest that PIKfyve inhibition reduces the ability of endocytosed cargo to reach terminal lysosomes, with no major effects on lysosomal proteolysis. Interestingly, like the vacuolation phenotype, apilimod mediated accumulation was exacerbated in *Inpp4b^-/-^* MEF, further indicating a role for Inpp4b in lysosomal dynamics (Bissig et al., 2017; Zhang et al., 2012).

### Inpp4b-deficiency exacerbates apilimod-mediated lysosome fusion-fission dynamics

Since apilimod blocks lysosomal accumulation of endocytic cargo such as LY, we adjusted our treatment conditions so that MEF were pulsed first with LY prior to short treatment with 10 nM and 500 nM apilimod doses for 1 h (**Fig. 2A**). LY revealed that individual lysosome size was similar in untreated *Inpp4b^+/+^* or *Inpp4b^-/-^* MEF, however apilimod induces lysosomal enlargement, where lysosomes in *Inpp4b^-/-^* MEF are significantly larger compared to *Inpp4b^+/+^* MEF (**Fig. 2A,B**). Accordingly, *Inpp4b^-/-^* MEF have reduced total lysosome numbers in untreated and apilimod-treated cells (**Fig. 2A,C**), however the relative LY^+^ lysosome content was unchanged upon apilimod treatment (**Fig. 2A,D**). These data conform to a model whereby apilimod treatment promotes lysosomal coalescence, which is manifested as increased individual lysosome volume and reduced total lysosome numbers (Choy et al., 2018; Saffi et al., 2019, 2021). Notably, *Inpp4b*-deficiency exacerbates apilimod-induced lysosomal coalescence, thus providing further evidence of a role for Inpp4b in lysosomal homeostasis and dynamics which is revealed upon apilimod treatment.

**Figure 2.**
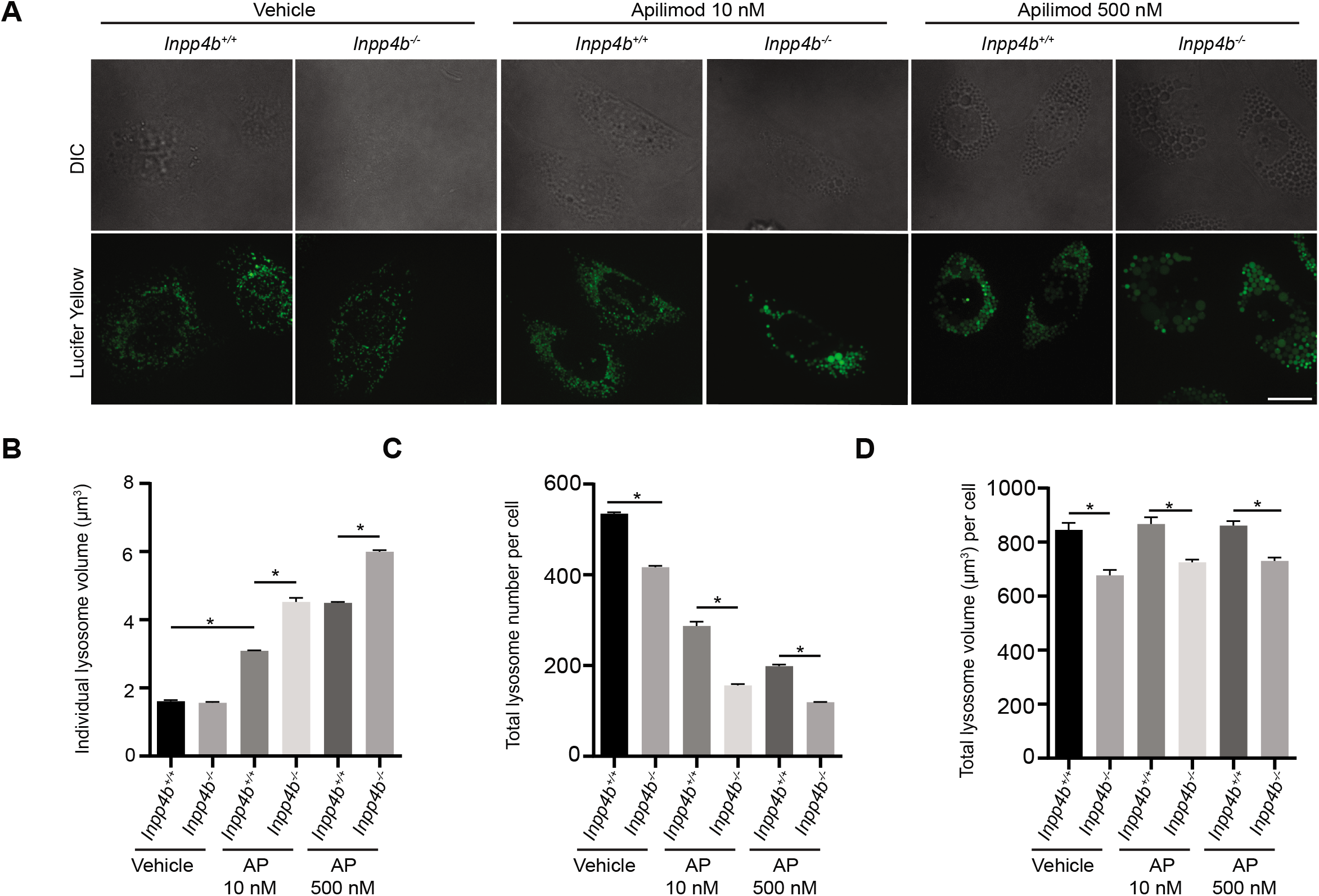
Lysosome volumetric and number analysis in Inpp4b and PIKfyve suppressed conditions. (**A**) Lysosomes of *Inpp4b^+/+^* or *Inpp4b^-/-^* MEFs pre-labelled with Lucifer Yellow and exposed to vehicle or apilimod at indicated concentrations for 1 h. Fluorescence micrographs represent Z-projections of 25-30 Z-planes acquired through spinning disc confocal microscopy. Image analysis performed for individual lysosome volume (**B**), lysosome number per cell (**C**), and total lysosome volume per cell (**D**). Scale bar: 25 μm. Data represent mean ± S.E.M. from three independent experiments, with 25-30 cells assessed per treatment condition per experiment. Significance measured through one way ANOVA and Tukey’s *post-hoc* represented as * in comparison to indicated conditions (p<0.05).

Inhibition of lysosomal fission has been proposed as an explanation for apilimod-induced lysosome enlargement (Choy et al., 2018; Saffi et al., 2019, 2021). To investigate the specific consequence of *Inpp4b*-deficiency on “kiss-and-run” and/or “fusion-and-fission” cycles, live-imaging of LY labelled lysosomes was recorded. *Inpp4b^+/+^* and *Inpp4b^-/-^* MEF treated with either vehicle or apilimod were monitored for up to 15 minutes and lysosomal fission events were tallied. Despite similar fission rates in vehicle-treated *Inpp4b^+/+^* and *Inpp4b^-/-^* MEF, apilimod significantly reduced lysosome fission rates in *Inpp4b^+/+^* MEF, and remarkably fission was nearly abrogated in *Inpp4b^-/-^* MEF (**Fig. 3A,B, Movie S1-4**). This suggests that the enhanced lysosomal coalescence observed in *Inpp4b^-/-^* MEF upon apilimod treatment is due to an exacerbated inhibition of lysosomal fission. These results suggest that *Inpp4b*-deficiency alters lysosome dynamics such that they are sensitized to the lysosomal fission-inhibiting effects of apilimod.

**Figure 3.**
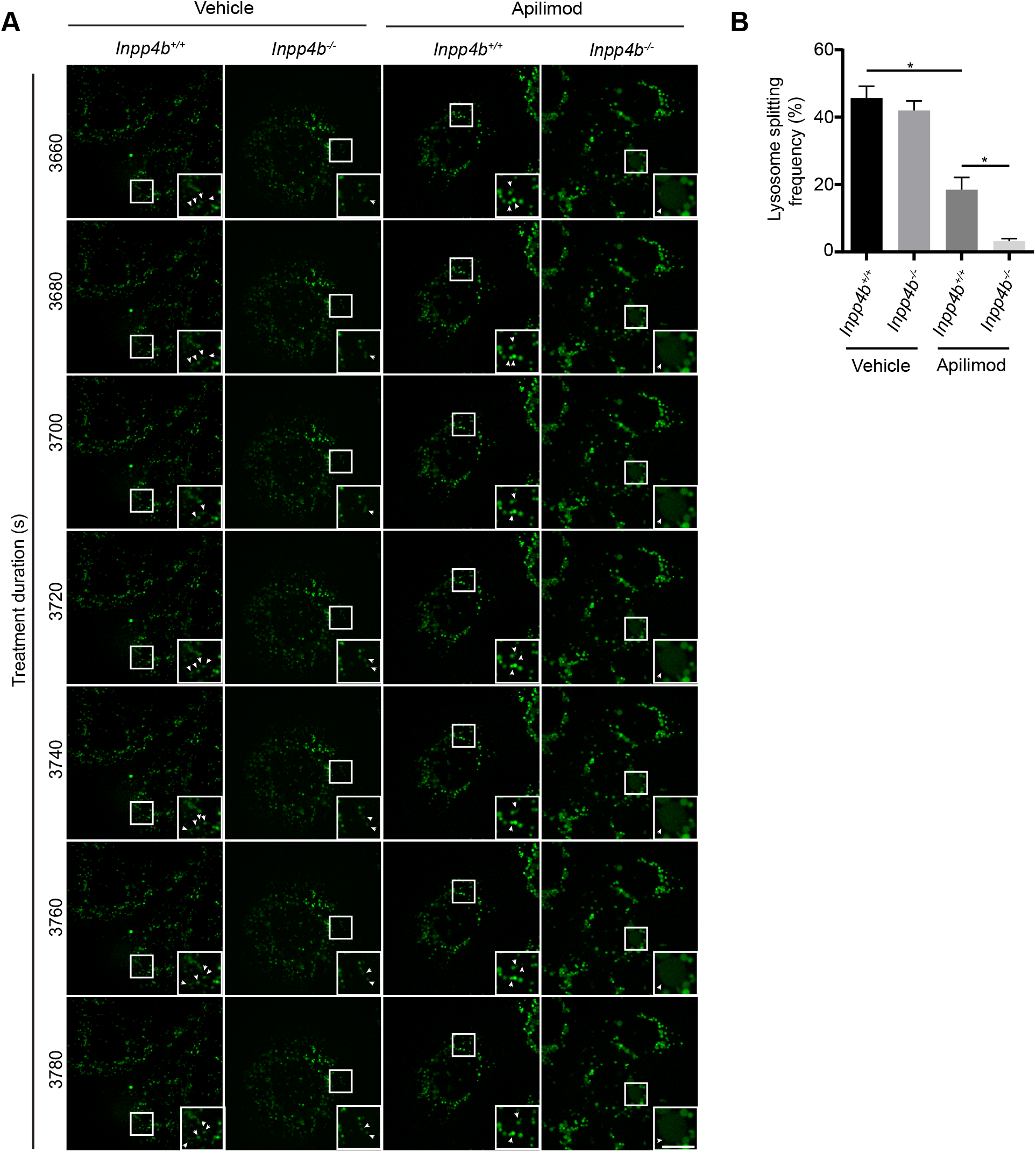
Live-cell imaging for lysosome “kiss-and-run” dynamics in INPP4B and PIKfyve suppressed cells. (**A**) *Inpp4b^+/+^* or *Inpp4b^-/-^* MEFs were pre-labelled with Lucifer Yellow followed by treatment with vehicle or 10 nM apilimod. Time in sec (s) refers to time post treatment. See movies 1 to 4 for full capture imaging videos. Inset represents an enlarged section from the field of view tracking individual lysosome particle dynamics over time. Scale bar: 10 μm. (**B**) Lysosome splitting frequency over 15 min for WT or Null MEFs treated with vehicle or apilimod demonstrate reduced splitting in PIKfyve inhibited WT MEFs and to a greater extent in PIKfyve inhibited Null MEFs, indicative of possible reduced lysosome fission. Data represent mean ± S.E.M. from three independent experiments. Significance measured through one way ANOVA and Tukey’s *post-hoc* represented as * in comparison to indicated conditions (p<0.05).

### Apilimod treatment induces differential effects on gene expression and lysosome function in Inpp4b^+/+^ and Inpp4b^-/-^ MEF

To shed light on the exacerbated phenotypes observed in *Inpp4b^-/-^* MEF treated with apilimod, we assessed the effects of apilimod treatment on lysosomal gene expression in *Inpp4b^+/+^* and *Inpp4b^-/-^* MEF at 48h. Evaluation of nuclear translocation of Transcription Factor EB (TFEB), a key regulator of lysosomal gene transcription (Sardiello et al., 2009), did not reveal any difference between *Inpp4b^+/+^* and *Inpp4b^-/-^* MEF with and without apilimod treatment (**Fig. 4A,B**). Next, transcript levels of lysosomal genes including *LAMP1, MCOLN1, CTSB, CTSD, ATP6V1D* and *ATP6V1H* were measured by TaqMan qPCR after 48 hours of vehicle or apilimod treatment. In vehicle treated cells, *Inpp4b^-/-^* MEF demonstrated modest, but significantly reduced expression of all lysosomal genes, except for *CTSD* (**Fig. 4C**). Strikingly, apilimod treatment led to a ~3-fold increase of all lysosomal transcripts tested in *Inpp4b^-/-^* MEF, whereas *Inpp4b^+/+^* MEF showed no significant changes under the same conditions (**Fig. 4C**). Immunoblotting of lysosomal proteins demonstrated that *Inpp4b^-/-^* MEF have reduced steady state expression of LAMP1, pre- and mature-cathepsin B, and V-ATPase V1H (**Fig. 4D,E**). After apilimod treatment, *Inpp4b^-/-^* MEF demonstrated significantly elevated expression of all lysosomal proteins tested, whereas *Inpp4b^+/+^* MEF only demonstrated elevated levels of the lysosomal membrane bound LAMP1 and V-ATPase V1H proteins (**Fig. 4D,E**). Next, to quantify differential effects of apilimod on lysosomal proteolytic function in *Inpp4b^+/+^* and *Inpp4b^-/-^* MEF, we used the membrane permeable cathepsin B substrate-Magic Red which upon hydrolysis forms membrane impermeable fluorescent cresyl-violet and accumulates within functional lysosomes (Bright et al., 2016). The results of this assay paralleled the transcript and protein expression levels of cathepsin B where apilimod treatment had no effect in *Inpp4b^+/+^* MEF. In *Inpp4b^-/-^* MEF, despite having a reduced steady state cathepsin B activity, apilimod treatment led to a more than 3-fold induction of cathepsin B activity (**Fig. 4F**). We speculate that the levels of transcription and translation of lysosomal genes, and the stability of lysosomal proteins may be differentially impacted by apilimod treatment in *Inpp4b^+/+^* and *Inpp4b^-/-^* MEF.

**Figure 4.**
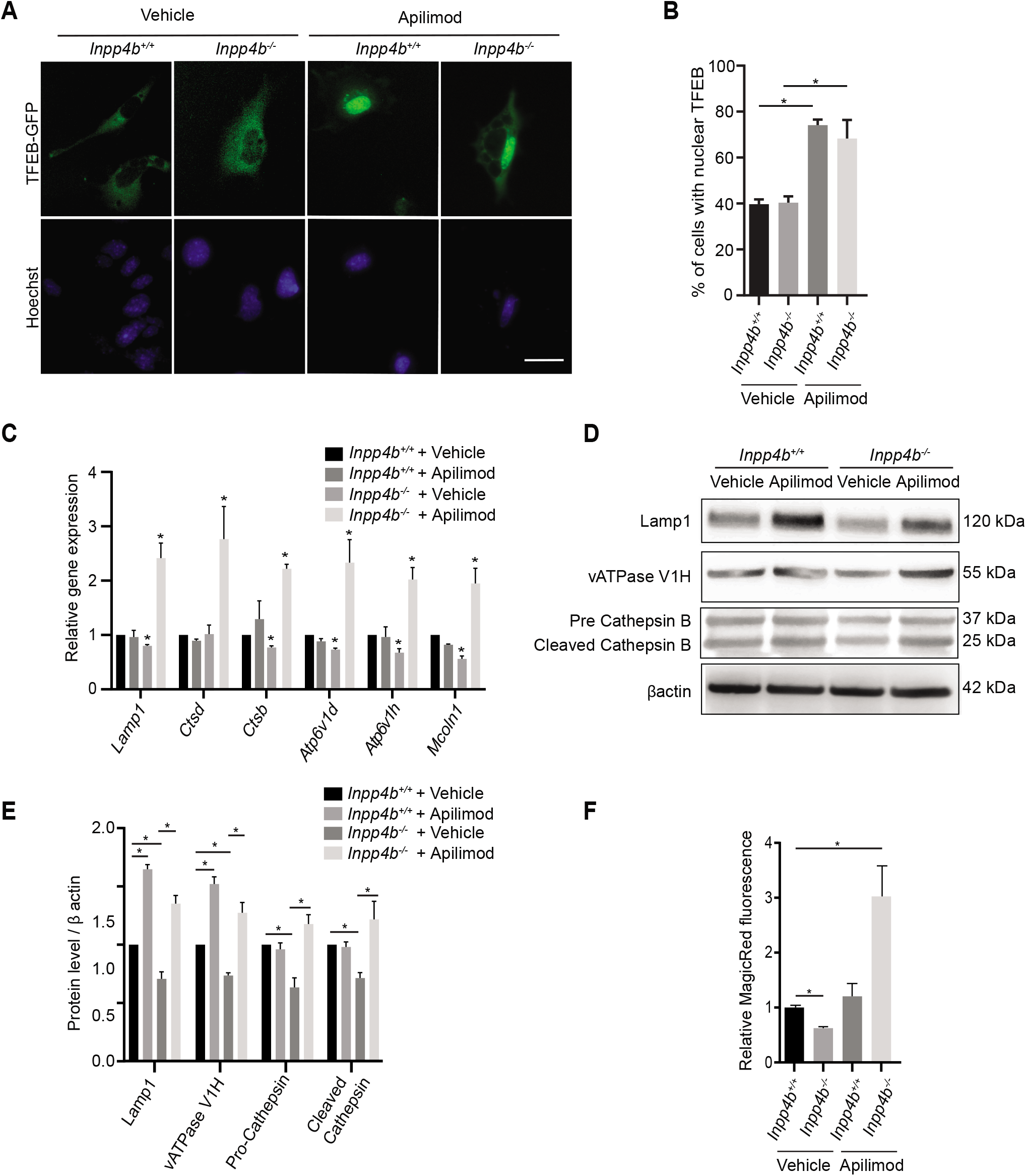
Inpp4b regulation of lysosome gene transcription and function. (**A**) *Inpp4b^+/+^* or *Inpp4b^-/-^* MEFs transiently expressing *pEGFP-TFEB* and treated with vehicle or apilimod 10 nM for 48 h. DNA was stained with Hoechst 33342. (**B**) TFEB nuclear translocation measured as percentage of cells demonstrating nuclear EGFP-TFEB for (**A**). Scale bar: 25 μm. (**C**) *Inpp4b^+/+^* or *Inpp4b^-/-^* MEFs treated with vehicle or apilimod 10 nM for 48 h followed by qRT-PCR analysis of lysosome gene expression normalized against Actb. Shown in mean ± SEM from three independent experiments. (**D**) Immunoblot of *Inpp4b^+/+^* or *Inpp4b^-/-^* MEFs treated with vehicle or apilimod 10 nM for 48 h followed by assessment of select lysosomal proteins as indicated. (**E**) Analysis of protein levels from (**D**) shown as mean ± SEM from three independent experiments. Data represent + SEM from 5 independent experiments with 30-35 cells assessed per treatment condition per experiment for *D-E.* (**F**) *Inpp4b^+/+^* or *Inpp4b^-/-^* MEF treated with vehicle or apilimod 10 nM for 48 h followed by Magic red incubation for 1 h. Quantification of Magic red intensity as measured through flow cytometry and mean ± SEM from three independent experiments. Significance measured through one way ANOVA and Tukey’s *post-hoc* represented as * in comparison to indicated conditions (p<0.05).

### Inhibition of autophagy by apilimod is potentiated in Inpp4b-deficient cells

PIKfyve inhibition has been demonstrated to inhibit autophagy in various physiological and pathological settings (Qiao et al., 2021; Martin et al., 2013; Hessvik et al., 2016). We evaluated how apilimod treatment affected autophagy in the context of *Inpp4b* deficiency. Firstly, we measured levels of MAP1LC3A/B (LC3), a membrane protein specifically expressed on autophagosomes by immunofluorescence imaging (Runwal et al., 2019). Total cellular levels of LC3 were similar in all vehicle-treated cells, and significantly elevated in *Inpp4b^+/+^* MEFs treated with apilimod, and further significantly elevated in apilimod-treated *Inpp4b^-/-^* MEF (**Fig. 5A,B**). Notably, the corresponding elevation in autophagosome levels induced by apilimod in *Inpp4b^-/-^* MEF was fully rescued by ectopic INPP4B expression (**Fig. S5A-C**). Autophagosome levels were further assessed by measuring LC-3 status by immunoblotting (Klionsky et al., 2016). During autophagy, cytosolic LC3-I (~16 kDa) is conjugated with phosphatidylethanolamine on the phagophore membrane to form LC3-II (~14 kDa). LC3-II is absent in *Inpp4b^+/+^* and *Inpp4b^-/-^* in vehicle-treated MEF (**Fig. 5D**), however apilimod treatment induced LC3-II levels in *Inpp4b^+/+^* MEF, and even further induced in *Inpp4b^-/-^* MEF (**Fig. 5D**). Similar observations were made in U2OS cells where INPP4B expression was silenced with siRNA (**Fig. S3E**). Autolysosome formation was assessed by colocalization LAMP1-mCherry fluorescence and LC-3 immunostaining. Apilimod treatment significantly increased the presence of autolysosome co-staining in *Inpp4b^+/+^* MEF, and *Inpp4b^-/-^* MEF showed a further significant increase (**Fig. 5A,C**).

**Figure 5.**
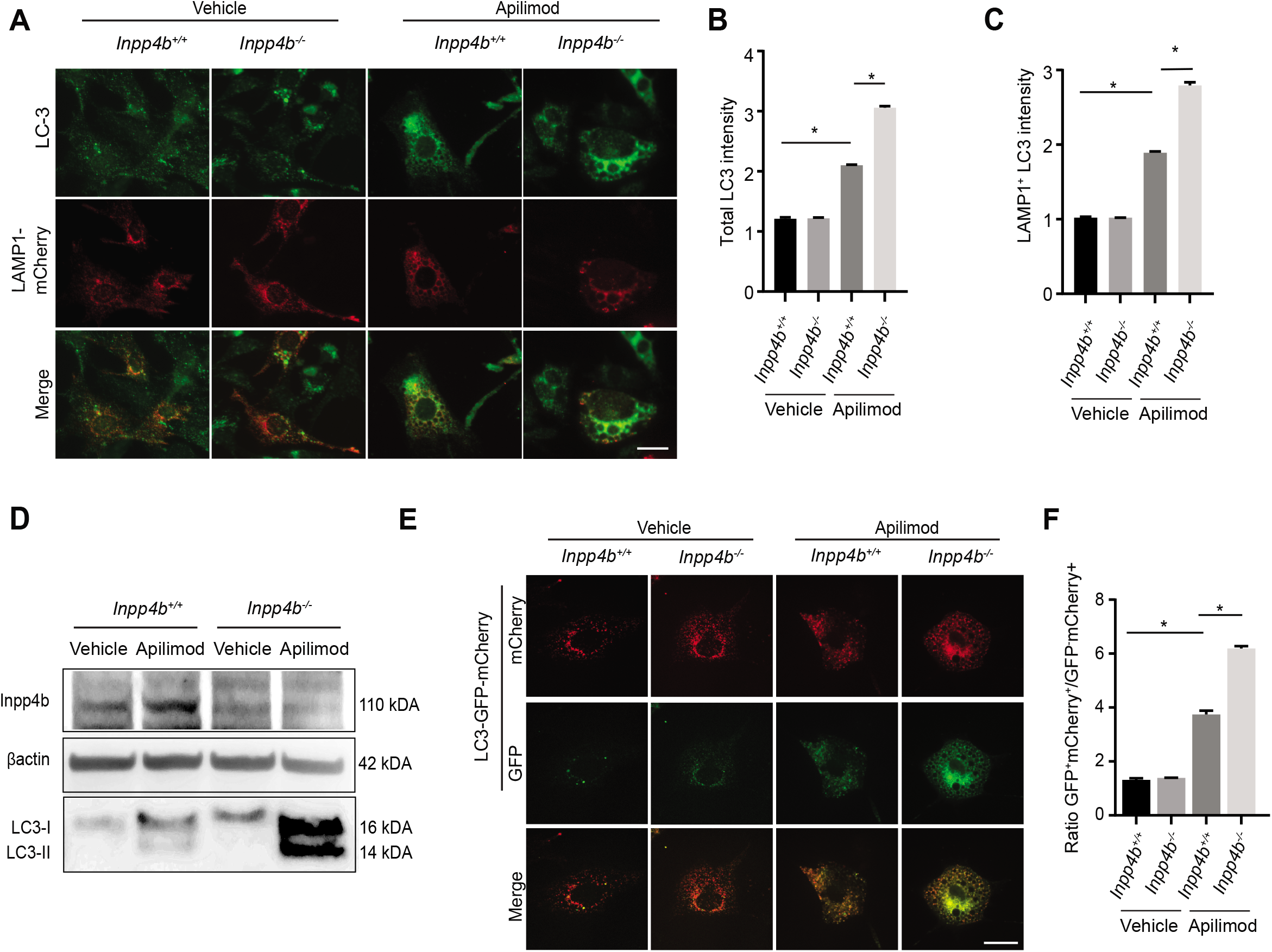
Autophagosome levels in response to Inpp4b and PIKfyve suppression. **(A)** *Inpp4b^+/+^* or *Inpp4b^-/-^* MEFs transiently expressing LAMP1-mCherry to mark lysosomes and treated with vehicle or apilimod 10 nM for 48 h, followed by immunostain against LC3. (**B**) Quantifications of LC3 puncta intensity per cell across indicated conditions to measure autophagosome levels and (**C**) LC3 intensity overlapping on LAMP1 positive lysosomes where increased LC-3/LAMP1 intensity ratio indicate autolysosome formation. (**D**) Immunoblot of *Inpp4b^+/+^* or *Inpp4b^-/-^* MEFs treated with vehicle or apilimod 10 nM for 48 h and assessed for protein levels of Inpp4b, LC3 and Beta actin as loading control. (**E**) *Inpp4b^+/+^* or *Inpp4b^-/-^* MEF transiently expressing *mCherry-EGFP-LC3B* and treated with vehicle or apilimod 10 nM for 48 h. (**F**) Quantification of LC-3 green puncta over red puncta intensity ratio. Scale bar: 25 μm. Data represent mean ± S.E.M. from three independent experiments, with 25-30 cells assessed per treatment condition per experiment for A-C and E-F. Significance measured through one way ANOVA and Tukey’s *post-hoc* represented as * in comparison to indicated conditions (p<0.05).

Finally, we estimated autophagic flux using a reporter system which expresses a fusion of LC3 protein, the acid-insensitive mCherry and the acid sensitive GFP (Zhang et al., 2013; N’Diaye et al., 2009). Quantification of the number of puncta (yellow or red) shows the total number of autophagosomes and autolysosomes and their ratio. Analysis of the spot number ratio of autolysosomes (mCherry^+^ GFP^-^) to autophagosomes (mCherry^+^ GFP^+^) allows the estimation of autophagosome to autolysosome transition (Sharma et al., 2019; Hansen and Johansen, 2011; Kim et al., 2020). In vehicle-treated MEF, we observed a dynamic switch from yellow fluorescence (mCherry^+^ GFP^+^) to red fluorescence (mCherry^+^ GFP^-^) indicating functional transition through autophagy in both *Inpp4b^+/+^* and *Inpp4b^-/-^* MEF (**Fig. 5E, F**). By contrast, apilimod blocks autophagy in MEF allowing us to estimate autophagic flux (**Fig. 5E-F**). These data demonstrate that *Inpp4b-*deficiency alone does not alter autophagic flux. These data demonstrate that *Inpp4b-*deficiency alone does not alter autophagic flux. However, apilimod treatment can reduce autophagic flux in MEF as observed in *Inpp4b^+/+^* MEF. As with other phenotypes, *Inpp4b^-/-^* MEF displays even further inhibition of autophagic flux.

### Inpp4b deficiency exacerbates apilimod-mediated PtdIns(3)P generation

Given the direct role of PIKfyve and Inpp4b on PtdIns metabolism, we assessed how PtdIns homeostasis may be differentially regulated in *Inpp4b^+/+^* and *Inpp4b^-/-^* MEF in response to apilimod-mediated PIKfyve inhibition. We first measured lysosomal PtdIns(3,4)P_2_ by immunofluorescence. As expected with *Inpp4b*-deficiency, greater total levels of PtdIns(3,4)P_2_ were observed (**Fig. 6A**). Similarly, ectopic LAMP1-mCherry expression allowed us to identify that there was significantly more lysosomal-associated PtdIns(3,4)P_2_ in *Inpp4b^-/-^* MEF compared to *Inpp4b^+/+^* MEF in the steady state (**Fig. 6A,C**). Apilimod treatment had no effect on PtdIns(3,4)P_2_ levels in *Inpp4b^+/+^* MEF, and there was a small but significant reduction in PtdIns(3,4)P_2_ levels compared to steady state levels (**Fig. 6A,C**). PtdIns(3)P levels were also measured by immunofluorescence in combination with ectopic LAMP1-mCherry expression. Unlike PtdIns(3,4)P_2_, total PtdIns(3)P levels and lysosomal PtdIns(3)P were unaffected by *Inpp4b* deficiency in untreated cells (**Fig. 6B**). However, apilimod treatment significantly induced PtdIns(3)P levels in both *Inpp4b^+/+^* and *Inpp4b^-/-^* MEF, where PtdIns(3)P induction was significantly more elevated in *Inpp4b^-/-^* MEF (**Fig. 6B,D**). Comparatively, examination of endosomal associated PtdIns(3)P as measured by GFP-2xFYVE^HRS^ puncta formation showed no difference in *Inpp4b^+/+^* and *Inpp4b^-/-^* MEF treated with apilimod (**Fig. S2B,D**). These observations are in agreement with previously conducted studies demonstrating no effect of apilimod on endosomal PtdIns(3)P (Chintaluri et al., 2018). (Chintaluri et al., 2018).

**Figure 6.**
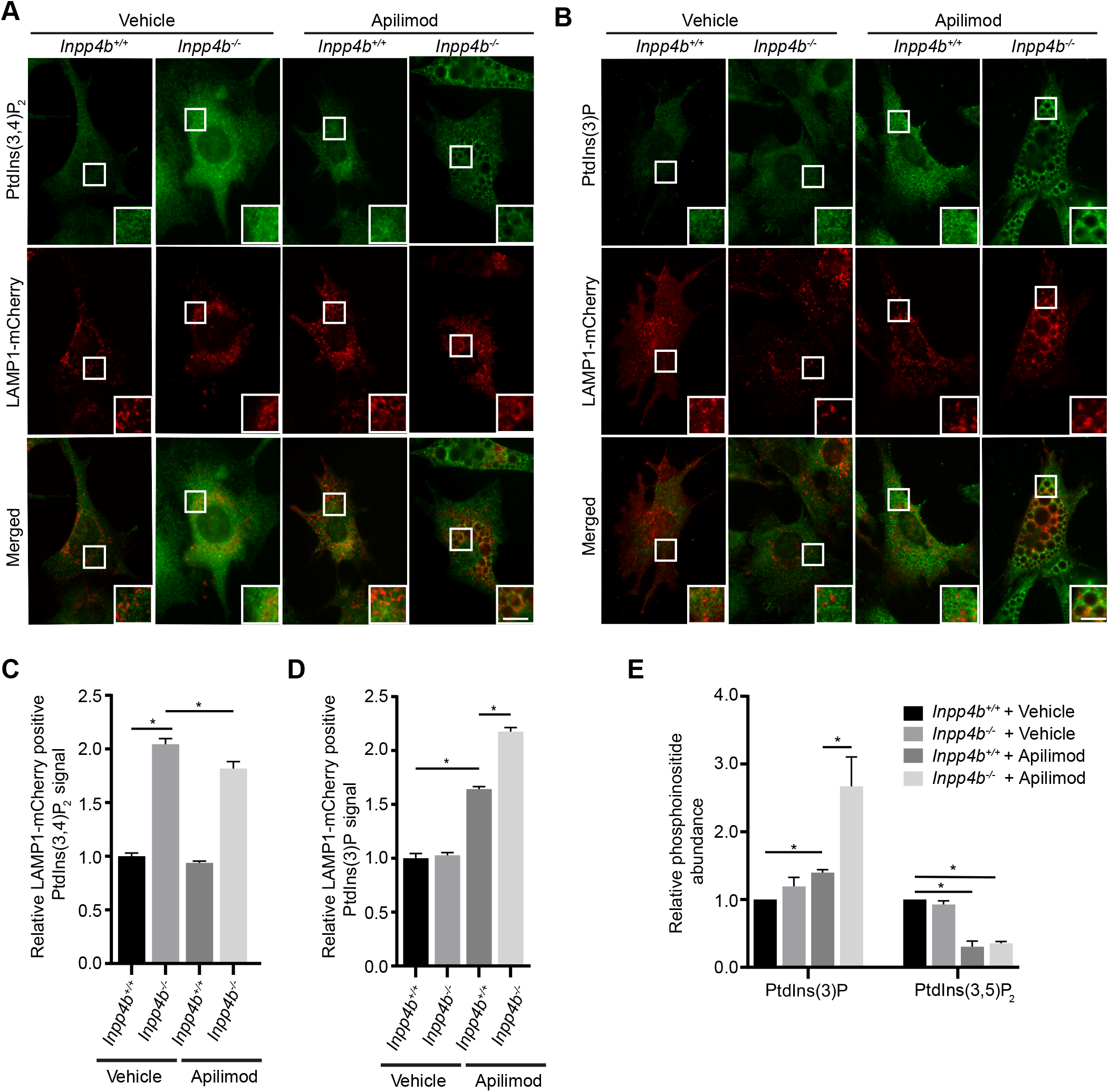
Phosphoinositide measurement in Inpp4b and PIKfyve suppressed cells. *Inpp4b^+/+^* or *Inpp4b^-/-^* MEFs transiently expressing LAMP1-mCherry and treated with vehicle or apilimod 10 nM for 48 h. (**A**) Total cell PI3,4P_2_ or (**B**) PI3P. Quantification of (**C**) PI3,4P_2_ fluorescence signal or (**D**) PI3P fluorescence signal respectively on LAMP1-mCherry positive regions within a cell. Scale bar: 5 μm. (**E**) ^3^H-*myo*-inositol incorporation and HPLC scintillation quantification of PI3P and PI3,5P_2_ for WT and Null MEFs treated with vehicle (Veh) or apilimod (AP). Data represent ± SEM from three independent experiments with 25-30 cells assessed per treatment condition per experiment for **A-D**. Significance measured through one way ANOVA and Tukey’s *post-hoc* represented as * in comparison to indicated conditions (p<0.05).

We also measured total cellular PtdIns abundance through ^3^H-*myo*-inositol labelling and HPLC scintillation and quantification. PtdIns(3,4)P_2_, was not detected in our experiments. PtdIns(3,5)P_2_ was readily detected and levels were not altered by *Inpp4b*-deficiency, but as expected apilimod treatment significantly reduced PtdIns(3,5)P_2_ levels in both *Inpp4b^+/+^* and *Inpp4b^-/-^* MEF (**Fig. 6E**). Similarly, total steady state PtdIns(3)P levels were unchanged in *Inpp4b^-/-^* MEF compared to *Inpp4b^+/+^* MEF. In comparison, apilimod treatment induced a modest increase of PtdIns(3)P in *Inpp4b^+/+^* MEF, but caused a striking ~3-fold increase in PtdIns(3)P in *Inpp4b^-/-^* MEFs (**Fig. 6E**). Altogether our analysis of PtdIns show that *Inpp4b*-deficient MEF possess elevated levels of PtdIns(3,4)P_2_, which was expected given that it is the main substrate of Inpp4b. However, *Inpp4b*-deficiency did not affect steady state PtdIns(3)P levels in MEF. Surprisingly, apilimod treatment led to a dramatic increase in total and lysosomal PtdIns(3)P in *Inpp4b^-/-^* MEF, suggesting a key role for Inpp4b in regulating PtdIns(3)P in the exacerbated lysosome enlargement phenotype.

### Apilimod-mediated lysosome enlargement is driven by VPS34-mediated PtdIns(3)P production

We sought to gain a further understanding of the role of PtdIns(3)P in mediating the exacerbated lysosomal enlargement upon apilimod treatment in *Inpp4b^-/-^* MEF. Bafilomycin-A1 (BafA1) has been previously reported to preclude the appearance of cytoplasmic vacuoles in apilimod treated cells by inhibiting apilimod-mediated PtdIns(3)P induction (Sbrissa et al., 2018). We observed little effect of BafA1 in apilimod treated *Inpp4b^+/+^* MEF. In contrast, generation of massively enlarged vacuoles observed in apilimod-treated *Inpp4b^-/-^* MEF was reversed with concurrent BafA1 treatment to levels similar to apilimod-*Inpp4b*^+/+^ MEFs (**Fig. S6A-C**) in a manner consistent with previous reports (Sbrissa et al., 2018), indicating a role for V-ATPase in the vacuolation phenotype.

Since the major source of cellular PtdIns(3)P is the class III PI3K-VPS34, we posited that depletion of cellular PtdIns(3)P may also attenuate generation of massively enlarged vacuoles in apilimod-treated *Inpp4b^-/-^* MEF. For these experiments, we treated MEF concurrently with apilimod and the specific VPS34 inhibitor, VPS34-IN1 (Munson et al., 2015; Bago et al., 2014). Firstly, PtdIns(3)P immunofluorescence performed on VPS34-IN1-treated MEF demonstrated that this inhibitor can effectively diminish total cellular PtdIns(3)P levels (**Fig. 7A**), and furthermore PtdIns(3)P immunofluorescence colocalization with mCherry-LAMP1 revealed that VPS34-IN1 can also significantly reduce levels of lysosome associated PtdIns(3)P (**Fig. 7A,B**). VPS34-IN1 had little effect on lysosome size and number in *Inpp4b^+/+^* MEF, possibly a result of the minimal changes in PtdIns(3)P levels in these cells. By contrast in *Inpp4b^-/-^* MEF, VPS34 inhibition effectively diminished the lysosomal enlargement to levels which are comparable to PIKfyve-inhibited *Inpp4b^+/+^* MEF (**Fig. 7C-E**). These results with VPS34-IN1 indicate that elevated levels of total and lysosomal PtdIns(3)P are VPS34-derived, and are critical to mediating the exacerbated lysosomal enlargement in *Inpp4b^-/-^* MEF upon apilimod treatment.

**Figure 7.**
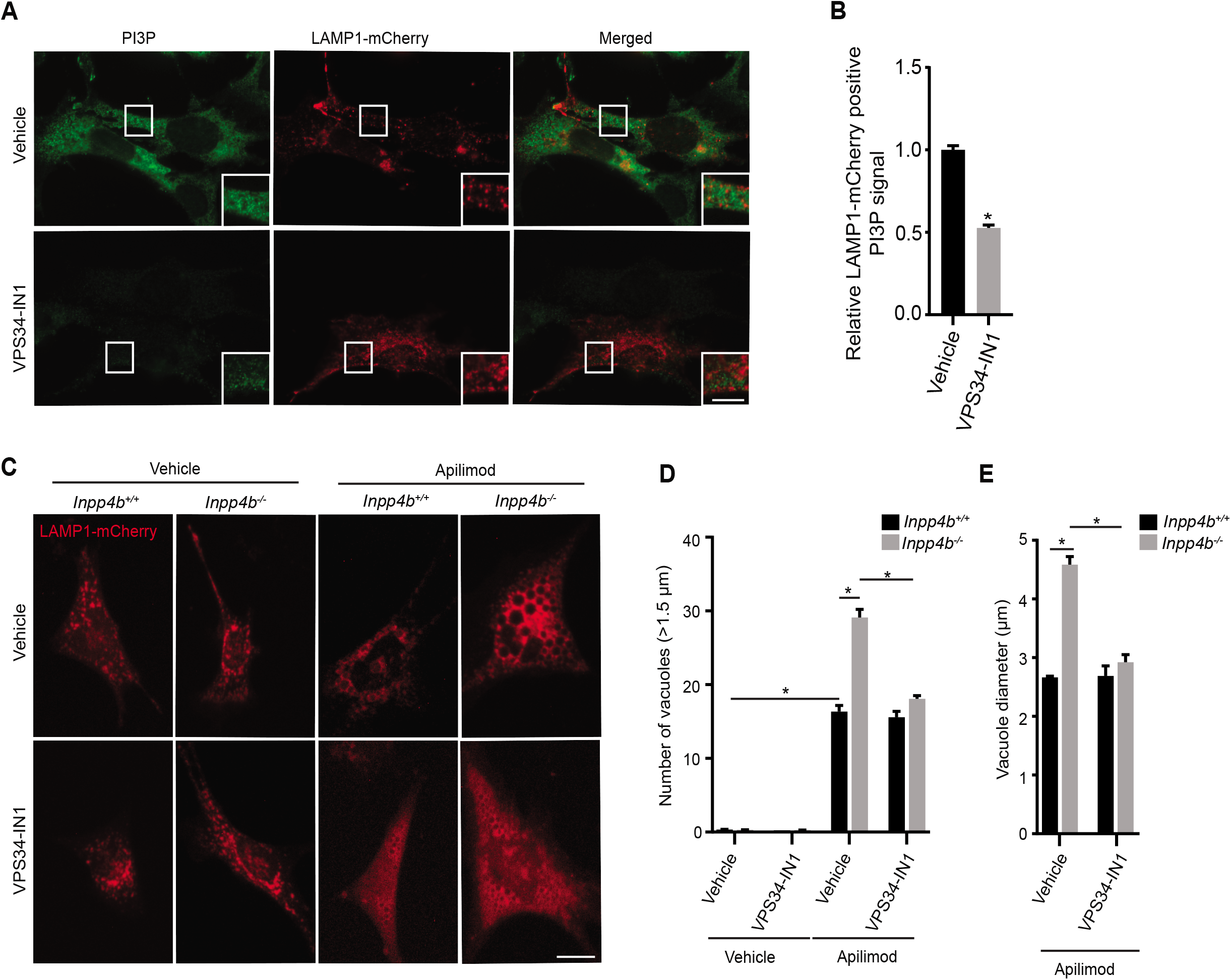
Effect of VPS34 and Inpp4b inhibition on lysosome morphology. (**A**) *Inpp4b^+/+^* MEF transiently expressing LAMP1-mCherry and treated with vehicle or 500 nM VPS34-IN1 for 48 h followed by PI3P immunostain. (**B**) Quantification from of PI3P fluorescence signal overlayed on LAMP1-mCherry positive regions within a cell. Scale bar: 5 μm. **(C)** WT or Null MEFs transiently expressing LAMP1-mCherry and treated with vehicle, apilimod 10 nM, or VPS34-IN1 500 nM at various combinations for 48 h. (**D**) Quantification of LAMP1 positive vacuoles greater than 1.5 μm in diameter per cell and **(E)** mean vacuole diameter (μm). Scale bar: 20 μm. Data represent ± SEM from three independent experiments with 25-30 cells assessed per treatment condition per experiment. Significance measured through one way ANOVA and Tukey’s *post-hoc* represented as * in comparison to indicated conditions (p<0.05).

## DISCUSSION

Lysosomes, the primary catabolic organelles in the cell, play pivotal roles in many cellular processes including cell differentiation, plasma membrane repair, programmed cell death, nutrient sensing and stress responses (Korolchuk et al., 2011). Thus, exquisite control of lysosome homeostasis, including the dynamic mechanisms that control total lysosomal content, number and size are critical to maintaining normal lysosomal and cellular functions. Emerging data indicates a role for PtdIns homeostasis in maintaining lysosome functions. For instance, inhibition of PIKfyve leads to disrupted lysosomal “fusion-fission” cycles by compromising “fission”, thereby generating enlarged coalescent lysosomes (Bissig et al., 2017; Choy et al., 2018; Saffi et al., 2021). PIKfyve inhibition also disrupts endocytic cargo delivery, autophagosome formation and autophagic flux, possibly a result of lysosome membrane perturbations where lysosomes fail to resolve from other lysosomes, late endosomes and autophagosomes (Bissig et al., 2017; Sharma et al., 2019). PIKfyve has been shown to functionally coordinate with the Class III PI3-Kinase VPS34 through its catalytic product PtdIns(3)P. PtdIns(3)P generated by VPS34 serves as a membrane localization target for PIKfyve, as well as a substrate-precursor for PtdIns(3,5)P_2_ synthesis (Ikonomov et al., 2015). Together the functions of PIKfyve and VPS34 recruit and regulate components of endosomal and lysosomal recycling (Giridharan et al., 2021). This PIKfyve-VPS34 crosstalk indicates the existence of other PIKfyve cross-talk with other PtdIns-modifying enzymes.

Recent studies suggest that the lipid-phosphatase INPP4B is a regulator of lysosome-associated functions. INPP4B depletion in triple negative breast cancer resulted in delayed EGFR trafficking from early endosomes to late endosomes/lysosomes (Liu et al., 2020). *INPP4B* expression in acute myeloid leukemia (AML) regulates lysosomal biogenesis and function which is critical for leukemia stem cell maintenance, differentiation and chemoresistance (Woolley et al., 2021). A role for INPP4B in lysosomal functions is further supported by the observation that elevated expression of INPP4B in *PIK3CA*-mutant ER^+^ breast cancer cells induce formation of late endosomes and lysosomes, increase cargo trafficking toward lysosomes, and promote endosomal sequestration and lysosomal degradation of key signaling proteins (Rodgers et al., 2021). Together, these emerging data support a role for INPP4B in the enhancement of function, and cellular content of lysosomes. Indeed, this study further supports this notion as *Inpp4b*-deficient MEF consistently demonstrated reduced total lysosomal content and lysosome numbers as measured by LAMP1 immunofluorescence (Fig. 1), lysotracker labelling (Fig. S1) and lysosomal accumulation of LY (Fig. 2 & 3) all observed in the absence of apilimod treatment. Furthermore, lysosomal gene (and protein) expression levels are suppressed in *Inpp4b^-/-^* MEF, together suggesting a role for Inpp4b in promoting and/or maintaining lysosomal content by controlling biogenesis through transcriptional mechanisms (Fig 4). Since INPP4B and PIKfyve both regulate lysosome function, we sought to shed light on interactions between the action of these two enzymes.

Apilimod-mediated PIKfyve inhibition in MEF leads to formation of numerous enlarged translucent cytoplasmic vacuoles readily seen by light microscopy (Fig 1A). The key findings made in this study stem from the observations that apilimod treatment of *Inpp4b^-/-^* MEF generates much larger cytoplasmic vacuoles compared to apilimod-treated *Inpp4b^+/+^* (Fig 1A). This was the first suggestion of crosstalk between INPP4B and PIKfyve. To gain further insight into the exacerbated phenotype in *Inpp4b*-deficient MEF, we first confirmed that enlarged vacuoles were of lysosomal origin as proposed by Choy and colleagues in both *Inpp4b^+/+^* and *Inpp4b^-/-^* MEF (Choy et al., 2018). Notably, treatment of MEF with apilimod prior to LY pulses revealed that apilimod inhibits lysosomal accumulation of cargo, a phenotype which which was exaggerated in *Inpp4b^-/-^* MEF (**Fig. S4**). Preloading of MEF with LY permitted effective lysosomal accumulation, which thereby enabled the clear visualization and quantitation of lysosomal features including size and number which were enhanced in *Inpp4b^-/-^* MEF. Furthermore, Inpp*4b^-/-^* MEF are exquisitely sensitized to apilimod, as demonstrated by the nearly abrogated fission, compared to the fission rates observed in *Inpp4b^+/+^* MEF. Altogether, our findings point to a role for Inpp4b in lysosomal dynamics, specifically *Inpp4b*-deficiency sensitizes cells to apilimod-mediated PIKfyve inhibition by disrupting lysosomal fission. Enhanced disruption of lysosomal fission provides an explanation for the enlarged vesicles observed in *Inpp4b^-/-^* MEF.

In our attempts to further understand the underlying biology responsible for the exaggerated response to apilimod observed in *Inpp4b^-/-^* MEF, lysosomal gene and protein expression was also measured after 48 hours of apilimod treatment. Expression of *Lamp1, Ctsd, Ctsb, Atp6v1d, Atp6v1h* and *Mcoln1* transcripts were unchanged in *Inpp4b^+/+^* MEF; by contrast *Inpp4b^-/-^* MEF demonstrated a significant 2-3 fold increase in the expression of each of these transcripts (Fig. 4). Nuclear localization of Tfeb was similar when *Inpp4b^+/+^* and *Inpp4b^-/-^* MEF were compared, suggesting that Inpp4b is not functioning through Tfeb. Protein expression provided a different picture; membrane bound proteins Lamp1 and V-ATPase (V1H) subunit were significantly induced upon apilimod treatment as would be expected by increased total lysosomal content suggesting post transcriptional stabilization or transcript or post-translational stabilization of protein.

Apilimod has previously been demonstrated to inhibit autophagy (Qiao et al., 2021; Martin et al., 2013; Hessvik et al., 2016; Sharma et al., 2019). Accordingly, we observed accumulation of LC3 puncta in MEF, many of which were LAMP1 positive indicating functional autolysosome formation. As an inhibitor of autophagy, apilimod revealed that *Inpp4b^-/-^* MEF have enhanced rates of autophagic flux. How the enhanced inhibition of lysosomal fission observed in apilimod-treated *Inpp4b^-/-^* MEF alters autolysosome formation is currently unknown but may reveal effects on autophagic flux. Furthermore, disruption of autophagy prevents turnover of lysosomal proteins (de Campos et al., 2020; Sharma et al., 2019), and may explain elevation of lysosomal proteins (LAMP1 and V-ATPase), despite the absence of transcriptional upregulation in *Inpp4b^+/+^* MEF. Apilimod treated *Inpp4b^-/-^* MEF on the other hand demonstrated elevated levels of cathepsin-B protein and activity in conjunction with autophagic flux inhibition. This combination of events may provide an explanation for increased sensitivity to apilimod observed in *Inpp4b^-/-^* MEF. It is currently unclear if elevated cathepsin-B protein and activity result due to disrupted feedback regulation due to apilimod effects.

Finally, we measured intracellular phosphoinositide levels in apilimod treated *Inpp4b^+/+^and Inpp4b^-/-^* MEF. As expected, *Inpp4b^-/-^* MEF cells display increased total and lysosomal PtdIns(3,4)P_2_ levels in the steady state. The role of PtdIns(3,4)P_2_ is relatively unexplored in lysosomal homeostasis, however our work (herein and in other studies) suggests it may be associated with repressed lysosomal biogenesis (Woolley et al., 2021). On the other hand, PtdIns(3)P levels were unchanged in vehicle-treated MEF, which suggests that Inpp4b may only be a minor contributor and other major mechanisms may exist to control PtdIns(3)P levels. As previously reported, apilimod leads to moderately elevated levels of PtdIns(3)P, which we also observed in *Inpp4b^+/+^MEF* (Sbrissa et al., 2018). Surprisingly, apilimod treatment in *Inpp4b^-/-^* MEF led to a dramatic increase in both total cellular and lysosomal levels of PtdIns(3)P. This observation is paradoxical given that *Inpp4b*-deficiency should generate less PtdIns(3)P, and suggests that Inpp4b may regulate PtdIns(3)P levels through an indirect mechanism. We observed that concurrent apilimod and VPS34-IN1 treatment alleviated the exacerbated lysosomal enlargement observed with *Inpp4b-*deficiency, suggesting a direct role for VPS34 in apilimod-mediated PtdIns(3)P induction.

In conclusion, our results indicate that *Inpp4b*-deficiency sensitizes cells and lysosomes to a plethora of effects of apilimod-mediated PIKfyve inhibition, the most obvious of which is lysosomal enlargement greater than that observed in *wild-type* cells, likely due to blocked fission and thus, enhanced lysosome coalescence. Similar exacerbated consequences are observed for various other phenotypes associated with PIKfyve inhibition including disrupted autophagy and reduced cargo delivery to lysosomes, (Bissig et al., 2017; Sharma et al., 2019; Dayam et al., 2015; Mironova et al., 2016). Importantly, we identify a novel paradoxical role for INPP4B in the regulating induction of PtdIns(3)P levels upon PIKfyve inhibition.

## MATERIALS AND METHODS

### MEF preparation, Cell culture conditions, Transfections, Drug treatment

Immortalized *Inpp4b^+/+^* and *Inpp4b^-/-^* MEF were generated as previously detailed (Mangialardi et al., 2019). MEF and U2OS cells were maintained in Dulbeco’s Modified Eagle Medium (DMEM) supplemented with 10% Fetal Bovine Serum (FBS). MEF were transiently transfected with *pTWIST-mCherry, pTWIST-Inpp4b-mCherry, mCherry-Lamp1, mCherry-EGFP-LC3B, pEGFP-2xFYVE, pEGFP, GFP-Inpp4b, GFP-Inpp4b (C845A)* and *pEGFP-TFEB.* U2OS cells were stably transfected with *mCherry-Lamp1* through selection with 200 μg/ml G418 for 10 days. Transfections of MEF and U2OS cells performed with Fugene HD (Promega, Madison, WI) at 3:1 of DNA:Fugene ratio for 24 h followed by washing and supplementation with complete DMEM growth media. siRNA-mediated gene silencing for *INPP4B* in U2OS cells carried out using DharmaFECT1 Transfection reagent (GE Dharmacon, Lafayette, CO). Briefly, 0.1 nmol of non-targeting or *Inpp4b* siRNA (GE Dharmacon) mixed with 2 μL of DharmaFECT1 Transfection reagent in DMEM media without FBS was added to U2OS cells for 24 h, followed by washing off the transfection mix with PBS and growth of cells for 48 h with treatment before imaging and western blot.

MEF and U2OS cells treated with apilimod or VPS34-IN1 (Selleck Chemicals, Houston, TX) to inhibit PIKfyve or VPS34 functions respectively, or bafilomycin A1 to inhibit lysosomal vacuolar ATPase and impair lysosome function (Sigma-Aldrich, Oakville, ON) at dose and durations as indicated.

### Retroviral transduction

3.0 x 10^6^ HEK 293T cells grown in a 10 cm dish for 24 followed by calcium phosphate transfection. Briefly, 10 μg of retroviral plasmid *pWZL hygro SV40 T-Large* mixed with 5 μg of pCL-Eco retroviral packaging vector and 2M CaCl_2_ to a final volume of 300 μL in sterile water. The transfection mix was supplemented with equal volume of 2X HEPES-Buffered saline (HBS; 140 mM NaCl, 1.5 mM Na_2_HPO_4_) followed by addition to the cells. Following 24 h post transfection, the media was changed and supplemented with complete DMEM growth media. Media was collected 48 h and 72 h post transfection. Virus enriched media was filtered through a 0.45-micron filter, supplemented with 8 μg/ml protamine sulfate and added to MEF cells grown in 10 cm dishes. Infections repeated every 8 h and MEF cells selected for 4 days with 75 μg/ml hygromycin B.

### Lysosome labelling

Lysosomes of MEF cells were labelled with 1 mg/ml Lucifer Yellow (LY; Thermo Fisher Scientific, Mississauga) for 2 h in complete growth media at 37 ^0^C and 5% CO_2_, followed by washing in phosphate-buffered saline (PBS) and supplementation of complete media for 1 h. Lysotracker red (Thermo Fisher) was also used to label lysosomes of MEF cells through incubation with cells at 1 μM for 30 min in complete growth media. Lysotracker red washed off with PBS and cells supplemented with complete growth media. Magic red (Abcam, Cambridge, UK) was used to assess lysosomal cathepsin B activity of MEF cells by incubation for 1 h according to manufacturer instructions followed by washing off with PBS and supplementation with complete growth media.

### Immunofluorescence

Immunolabeling of cells following apilimod treatment was performed by fixation with 4% (v/v) paraformaldehyde for 15 min, permeabilization with 100% ice-cold methanol for 5 min, and blocking in 3% BSA (v/v) in PBS. Cells incubated with rabbit monoclonal antibody against mouse LC3B (1:200; Cell Signaling) and Alexa Fluor 488-conjugated goat polyclonal antibody against rabbit IgG (1:1000; Thermo Fisher). Alternatively, immunostaining performed with rat monoclonal antibody against mouse LAMP1 (1:200, Clone 1D4B; Thermo Fisher) and Dylight 488-conjugated donkey polyclonal antibody against rat IgG (1:1000; Bethyl, Montgomery, TX). Immunofixation performed with rabbit polyclonal antibody against mouse EEA1 (1:200; Cell Signaling) and Alexa Fluor 488-conjugated goat polyclonal antibody against rabbit IgG (1:1000; Thermo Fisher) according to manufacturer instructions. Total cell PI3,4P_2_ or PI3P immunofluorescence performed by fixation with 4% (v/v) paraformaldehyde for 15 min, permeabilization with 20 μM digitonin (Promega) in buffer A (20 mM PIPES pH 6.8, 137 mM NaCl, 2.7 mM KCl) for 30 min, and blocking with buffer A containing 5% normal goat serum and 50 mM NH_4_Cl. Immunostaining performed with Anti-PI3,4P_2_ IgG or Anti-PI3P IgG (Echelon, Salt Lake City, UT) and Dylight 488-conjugated Goat polyclonal antibody against mouse IgG (1:1000; Bethyl). Samples mounted onto microscope slides through DAKO fluorescent mounting media and imaged.

### Live- and fixed-cell microscopy

Manual quantifications of vacuole and lysosome size, number, LAMP1 positive vesicles, and TFEB-GFP localization were performed using EVOS-FL fluorescent inverted microscope controlled by EVOS XL core imaging system at 20x 0.4 N.A. (Thermo Fisher) objective. Spinning disc confocal microscopy was used to perform live imaging through Olympus IX81 inverted microscope connected to Hamamatsu C9100-13 EMCCD camera with 60x 1.35 N.A. objective and controlled by Volocity 6.3.0 (PerkinElmer, Bolton, ON). Time lapse live imaging performed with an environmental chamber set to 37^0^C and 5% CO_2_ in DMEM complete media. Fixed cells observed through ZEISS AxioImager M2 Epifluorescence microscope connected to AxioCam MRm CCD camera and controlled by AxioVision Software Version 4.8 at 20x 0.8 N.A. or 40x 1.4 N.A. objective (Carl Zeiss, Oberkochen, Germany).

### Image Analysis

To quantify lysosomes as enlarged vacuoles, LAMP1 positive vesicles were defined to have diameter greater than 1.5 μm followed by employing line plots to measure vacuole diameter using ImageJ. To measure the percentage of cells with nuclear TFEB, cells were scored to have nuclear TFEB if the nucleus had greater intensity compared to cytosol using ImageJ. To measure the percentage of lysosomes filled with LY per cell, images imported into ImageJ and number of LAMP1 positive vesicles were manually scored for presence of LY signal within the lysosome lumen.

To quantify LAMP1 or LC3 immunostaining, and GFP-2xFYVE fluorescence through ImageJ, intensity thresholding was applied to identify fluorescent structures and the mean intensity was obtained for each cell. To quantify LC3 intensity over LAMP1 positive structures, ImageJ used to threshold for LAMP1-mCherry signal and generating a mask, which was applied to the green (LC3 immunostain) channel to measure LC3 intensity on LAMP1-mCherry positive regions. For MEF cells transiently expressing mCherry-eGFP-LC3B, similar approach was used to determine LC3 green puncta intensity over LC3 red puncta structures where intensity ratio greater than 1 indicate formation of autolysosomes due to reduced autophagic flux. Similar image analysis technique was applied for MEF cells transiently expressing LAMP1-mCherry to evaluate PtdIns(3,4)P_2_ or PtdIns(3)P levels overlayed on LAMP1-mCherry positive regions within a cell.

To measure lysosome volume and number per cell, particle detection and volumetric tools from Volocity 6.3.0 were used. Briefly, Z-stack images imported into Volocity and punctate lysosome structures identified by applying a 2x cytosol intensity threshold to exclude cytosol and background. Further criteria to include particles greater than 0.3 μm^3^ removed noise-derived particles. Each cell was isolated by drawing region interest for individual cell analysis. Quantification of lysosome splitting frequency was performed through Imaris (BitPlane, Concord, MA) using ‘ImarisTrackLineage’ module, where lysosome splitting was defined as frequency of events where two particles were produced from a single particle.

### Western Blot

Whole cell lysates generated using 1X RIPA buffer supplemented with protease inhibitor. Proteins immunoblotted with the antibodies anti-LC3B (#3868), beta actin (#4967), INPP4B (#14543), LAMP1 (#3243) from Cell Signaling, anti-vATPase V1H (sc-166227) and cathepsin B (sc-365558) from Santa Cruz (Dallas, TX), or anti-INPP4b Neuromab clone N171/17 (Davis, CA).

### Flow cytometry

MEF cells were incubated with 10 μg/ml DQ-BSA (Invitrogen, Burlington, ON) or 1 mg/ml LY (Thermo Fisher) for 1 h to 6 h at 37 ^0^C, or Lysotracker red 1 μM for 30 min, or Magic red for 1 h. Alternatively, LAMP1-mCherry signal of U2OS cells recorded through flow cytometry following apilimod treatment. Briefly, cells washed twice with PBS at each time point and whole cell fluorescence recorded with the Beckman Coulter Cytoflex flow cytometer (Beckman, Brea, CO). A total of 10,000 events counted per condition per sample using the fluorescein isothiocyanate (FITC-A) channel for DQ-BSA and LY, or phycoerythrin (PE-A) channel for lysotracker red and LAMP1-mCherry and Magic red. Background signal was determined from non-labelled cells at time 0.

### Phosphoinositide labelling with ^3^H-myo-inositol and HPLC-coupled flow scintillation

MEF cells were incubated for two 24 h cycles with inositol-free media (MP Biomedical, CA), 10% dialyzed FBS (Gibco), 4 mM L-glutamine (Sigma Aldrich), 1x insulin-transferrin-selenium-ethanolamine (Gibco), 20 μCi/ml myo-[2-^3^-H(N)] inositol (PerkinElmer, MA) and indicated treatment conditions. Cells were washed twice with 1x PBS between each 24 h cycle. Lipid precipitation induced, followed by lipid deacylation, extraction and phosphoinositide separation by HPLC (Agilent Technologies, Mississauga, ON) through anion-exchange 4.6 x 250-mm column (Phenomenex, Torrance, CA) as previously mentioned (CH et al., 2018). β-RAM 4 (LabLogic, Brandon, FL) and 1:2 ratio of eluate to scintillation fluid (LabLogic) was used to detect radiolabeled eluate, followed by analysis with Laura 4 software (Ho et al., 2016)..

### Quantitative RT PCR

RNA isolation from MEF cells performed through Qiagen RNeasy mini kit (Qiagen, Venlo, Netherlands). Superscript IV Vilo cDNA synthesis kit (Thermo Fisher) was used to reverse transcribe equal amount of mRNA. The resulting cDNA was amplified through quantitative PCR using TaqMan Fast Advanced Master mix (Applied Biosystems, Foster City, CA) according to manufacturer instructions in presence of Taqman assays with QuantStudio 3 Real-Time PCR system (Thermo Fisher) controlled by QuantStudio Design and Analysis Software version 1.2 (Thermo Fisher). Taqman assays include Actb (Mm02619580_g1), CtsD (Mm00515586_m1), CtsB (Mm00514443_g1), Atp6v1h (Mm01224350_m1), Atp6v1d (Hs00211133_m1), Lamp1 (Mm01217068_g1) and Mcoln1 (Mm01211241_g1) and were performed in triplicates. Relative quantification (ΔΔCt method) was used to determine gene expression normalized to Actb and vehicle treated WT MEF.

### Statistical Analysis

All experiments conducted independently at least three times and statistical analysis to compare significance between multiple conditions performed through one-way ANOVA test coupled with Tukey’s *post hoc* test. P values less than 0.05 were statistically significant.

## ACKNOWLEDGEMENTS

We thank all the members of the Salmena lab for thoughtful discussions and constructive criticism. The following plasmids were obtained from Addgene (Cambridge, MA): mCherry-Lysosomes-20 was a gift from Michael Davidson (Addgene plasmid # 55073), pEGFP-2xFYVE was a gift from Herald Stenmark (Addgene plasmid # 140047), pBABE-puro mCherry-EGFP-LC3B was a gift from Jayanta Debnath (Addgene plasmid # 22418), pEGFP-N1-TFEB was a gift from Shawn Ferguson (Addgene plasmid # 38119).

## DATA AVAILABILITY

Upon Request

## AUTHOR CONTRIBUTIONS

GTS: Conceptualization, Data curation, Formal analysis, Investigation, Project Management, Methodology, Software, Visualization, Writing. EMM and JV: Resources. RJB: Resources, Writing

LS: Resources, Supervision, Funding Acquisition, Investigating, Project Management, Writing

## SUPPLEMENTAL FIGURE LEGENDS

**Supplementary Figure S1. Vacuole formation and rescue in Apilimod treated *Inpp4b^-/-^* MEFs. (A-C)** *Inpp4b^+/+^* or *Inpp4b^-/-^* MEFs treated with vehicle or apilimod 10 nM for 48 h, followed by Lysotracker red staining to monitor lysosome levels in Apilimod treated cells. (**D)** *Inpp4b^+/+^* or *Inpp4b^-/-^* MEFs transiently expressing LAMP1-mCherry to mark lysosomes followed by vehicle or 10 nM apilimod treatment for 48 h. Quantification of **(E)** Number of vacuoles (>1.5 μm in diameter) per cell across indicated treatments, and **(F)** Mean vacuole diameter (μm). Scale Bar: 25 μm. **(G)** *Inpp4b^-/-^*MEFs transiently expressing *pEGFP, GFP-Inpp4b, or GFP-Inpp4b (C845A)* and treated with vehicle or apilimod 10 nM for 48 h. Transfected cells indicated by white arrow vs. non-transfected cell(s) indicated by black arrow. Scale bar: 25 μm. Quantification of **(H)** Number of vacuoles (>1.5 μm in diameter) per cell across indicated treatments, and **(I)** Mean vacuole diameter (μm). Data represent ± SEM from three independent experiments with 25-30 cells assessed per treatment condition per experiment for *D-I* or 10,000 events per treatment condition per experiment for *B*. Significance measured through one way ANOVA and Tukey’s *post-hoc* represented as * in comparison to indicated conditions (p<0.05).

**Supplemental Figure S2. Vacuoles formed from Apilimod treated *Inpp4b^-/-^* MEFs are absent of endosomal markers. (A)** *Inpp4b^+/+^* or *Inpp4b^-/-^* MEFs transiently expressing LAMP1-mCherry and treated with vehicle or 10 nM apilimod for 48 h and immunostained for EEA1 endosomal marker. **(B**) WT or Null MEFs transiently expressing GFP-2x-fyve endosomal maker treated with vehicle or 10 nM apilimod for 48 h. Arrows indicate vacuoles absent of GFP-2x-fyve. **(C)** Quantification of percentage of LAMP1 vesicles positive for EEA1 endosomal marker from **(A). (D)** Quantification of GFP-2x-fyve puncta intensity from **(B).** Scale bar: 20 μm. Data represent ± SEM from three independent experiments with 25-30 cells assessed per treatment condition per experiment. Significance measured through one way ANOVA and Tukey’s *post-hoc* represented as * in comparison to indicated conditions (p<0.05).

**Supplementary Figure S3. Effect of INPP4B and PIKfyve suppression on lysosomes and autophagosomes in U2OS cells. (A)** U2OS cells stably expressing mCherry-LAMP1 and treated with vehicle or apilimod 10 nM for 48 h in control (siCtrl) and INPP4B-silenced cells (siINPP4B). **(B)** Quantification of mCherry-LAMP1 positive vacuole number (>1.5 μm in diameter) per cell across indicated treatments, **(C)** Mean vacuole diameter (μm) positive for mCherry-LAMP1 per cell across apilimod treated conditions and **(D)** mCherry-LAMP1 intensity per cell across indicated conditions as measured through flow cytometry. Scale bar: 20 μm. *(E)* U2OS cells from *(A)* immunoblotted for INPP4B, LC-3 and beta actin. Data represent ± SEM from three independent experiments with 40-50 cells assessed per treatment condition per experiment for *A-C* or 10,000 events per treatment condition per experiment for *D*. Significance measured through one way ANOVA and Tukey’s *post-hoc* represented as * in comparison to indicated conditions (p<0.05).

**Supplementary Figure S4. Endocytic function in Inpp4b and PIKfyve suppressed cells. (A)** *Inpp4b^+/+^* or *Inpp4b^-/-^* MEFs treated with vehicle or apilimod 10 nM for 48 followed by pulsing with Lucifer Yellow for 0, 1, 2 and 4 h. Quantification of Lucifer Yellow intensity as measured through flow cytometry. (**B**) *Inpp4b^+/+^* or *Inpp4b^-/-^* MEFs transiently expressing mCherry-LAMP1 and treated with vehicle or apilimod 10 nM for 48 h. Lucifer yellow pulsed for 0, 1, 2 and 4 h. **(C)** Quantification of percentage of mCherry-LAMP1 positive vesicles filled with Lucifer Yellow. Scale bar: 25 μm. (**D**) *Inpp4b^+/+^* or *Inpp4b^-/-^* MEFs treated with vehicle or apilimod 10 nM for 48 followed by pulsing with DQ-BSA for 0, 1, 2 and 4 h. Quantification of DQ-BSA intensity as measured through flow cytometry. Data represent ± SEM from three independent experiments with 25-30 cells assessed per treatment condition per experiment for *B-C* or 10,000 events per treatment condition per experiment for **A *and* D.** Significance measured through one way ANOVA and Tukey’s *post-hoc* represented as * in comparison to indicated conditions (p<0.05).

**Supplementary Figure S5. INPP4B expression rescue autophagosome defect from PIKfyve suppression. (A)** *Inpp4b^+/+^* or *Inpp4b^-/-^* MEFs transiently expressing mCherry or INPP4B-mCherry and treated with vehicle or apilimod 10 nM for 48 h. **(B)** Inset represents a single cell from **(A)** expanded field of view. **(C)** Comparison between non-transfected and transfected cells for LC3 puncta intensity per cell across indicated conditions. Scale bar: 20 μm. Data represent ± SEM from three independent experiments with 25-30 cells assessed per treatment condition per experiment. Significance measured through one way ANOVA and Tukey’s *post-hoc* represented as * in comparison to indicated conditions (p<0.05).

**Supplemental Figure S6. Bafilomycin-A1 reverses generation of massively enlarged vacuoles observed in apilimod-treated *Inpp4b^+/+^* MEF. (A)** *Inpp4b^-/-^* or *Inpp4b^-/-^* MEFs transiently expressing LAMP1-mCherry to mark lysosomes followed by vehicle or 10 nM apilimod treatment for 48 h. **(B)** Quantification of Number of vacuoles (>1.5 μm in diameter) per cell across indicated treatments, and **(C)** Mean vacuole diameter (μm). Scale Bar: 25 μm. Data represent ± SEM from three independent experiments with 25-30 cells assessed per treatment condition per experiment. Significance measured through one way ANOVA and Tukey’s *post-hoc* represented as * in comparison to indicated conditions (p<0.05).

**Supplementary Movie 1: Lysosome dynamics for vehicle treated *Inpp4b^+/+^* MEF cells.** Live cell imaging performed for WT MEF cells pre-labelled with Lucifer yellow, treated with vehicle and single z-plane images acquired every 20 sec for 15 min. Indicated are scale and time.

**Supplementary Movie 2: Lysosome dynamics for apilimod treated *Inpp4b^+/+^* MEF cells.** Live cell imaging performed for WT MEF cells pre-labelled with Lucifer yellow, treated with apilimod 10 nM for 1 h and single z-plane images acquired every 20 sec for 15 min. Indicated are scale and time.

**Supplementary Movie 3: Lysosome dynamics for vehicle treated *Inpp4b^-/-^* MEF cells.** Live cell imaging performed for Null MEF cells pre-labelled with Lucifer yellow, treated with vehicle and single z-plane images acquired every 20 sec for 15 min. Indicated are scale and time.

**Supplementary Movie 4: Lysosome dynamics for apilimod treated *Inpp4b^-/-^* MEF cells.** Live cell imaging performed for Null MEF cells pre-labelled with Lucifer yellow, treated with apilimod 10 nM for 1 h and single z-plane images acquired every 20 sec for 15 min. Indicated are scale and time.

